# Differential Regulation of NHE3 Expression in Type 1 and Type 2 Diabetic Intestine: Impaired Endosomal Regulation of NHE3 Expression in Type 1 Diabetes

**DOI:** 10.1101/2025.09.10.675364

**Authors:** Varsha Singh, Ruxian Lin, Laxmi Sunuwar, Jianbo Yang, Mark Donowitz, Rafiquel Sarker

**Author notes:** Correspondence: Varsha Singh Ph.D Johns Hopkins University, Departments of Medicine, Division of Gastroenterology and Hepatology Ross 933, 720 Rutland Ave, Baltimore, MD 21205, United States.

## Abstract

Chronic diarrhea is a frequent gastrointestinal complication in both type 1 (T1D) and type 2 diabetes (T2D), though the underlying mechanisms differ: T1D is linked to autonomic neuropathy and disrupted transporter regulation, while T2D is often linked to medications and intestinal inflammation. Using streptozotocin-induced mouse models of T1D and T2D, we observed increased luminal fluid in the small intestine of both. Given the role of Na^+^/H^+^ exchanger 3 (NHE3) in fluid absorption and its loss in most diarrheal diseases, we examined NHE3 expression across intestinal segments. In T1D, NHE3 protein was significantly reduced in the duodenum and jejunum without changes in mRNA, suggesting post-transcriptional regulation. In contrast, T2D mice exhibited reduced NHE3 protein and mRNA, restricted to the proximal colon. To investigate mechanisms underlying NHE3 loss in T1D, we evaluated endosomal scaffolding proteins involved in NHE3 trafficking. While our previous work showed that the Sorting Nexin-27 (SNX27)-retromer complex does not regulate NHE3 protein stability, we found that SNX17 was significantly decreased in the small intestine of T1D mice but unchanged in T2D. SNX17 knockdown in SK-CO15 cells reduced NHE3 activity and stability. A GST pull-down assay showed that SNX17 interacts with the C-terminus of NHE3. Mutation of the NHE3 distal NPxY motif disrupted this interaction, leading to reduced NHE3 expression and increased degradation. These findings reveal segment-specific and mechanistically distinct causes of diabetic diarrhea in T1D versus T2D, and identify SNX17 loss as a contributor to reduced NHE3 stability and activity in T1D, likely promoting diabetic diarrhea.

**New and Noteworthy:** This study identifies distinct mechanisms of impaired sodium absorption contributing to diabetic diarrhea in type-1 and type-2 diabetes. We identify SNX17 as a novel regulator of NHE3 in the small intestine, showing that SNX17 loss in T1D contributes to post-translational NHE3 destabilization. In contrast, T2D-associated NHE3 downregulation is transcriptional and confined to the colon. These findings reveal disease- and region-specific regulatory pathways that drive impaired fluid absorption in diabetes, with direct implications for the development of targeted therapies.

**Schematic overview of differential NHE3 regulation in diabetic diarrhea.:** In Type 1 Diabetes (T1D), reduced SNX17 expression in the small intestine causes increased degradation of NHE3 and impaired sodium absorption. In Type 2 Diabetes (T2D), SNX17 levels remain unchanged in the small intestine, but NHE3 expression is transcriptionally downregulated in the proximal colon. These distinct mechanisms contribute to segment-specific differential regulation of NHE3 in T1 and T2D diabetes.

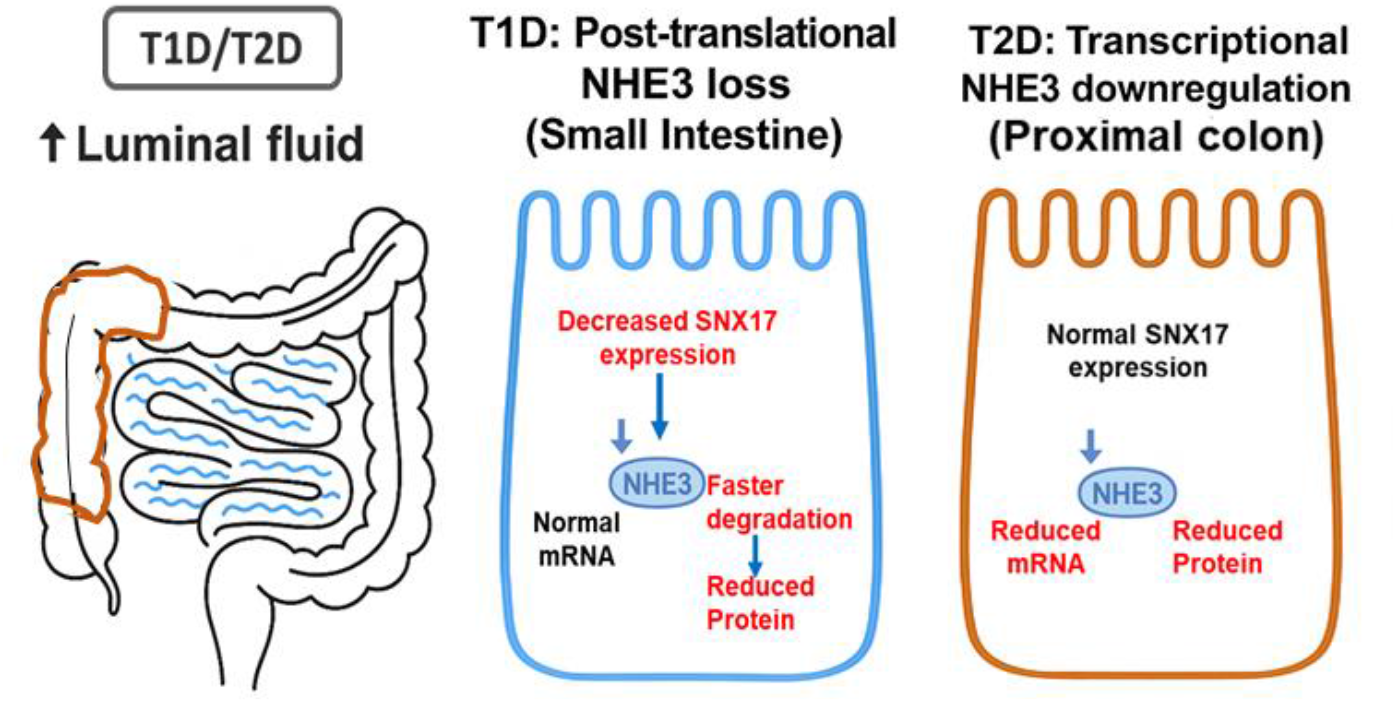

## Introduction

Diarrhea is a common and often underrecognized gastrointestinal complication in both type 1 diabetes (T1D) and type 2 diabetes (T2D). Overall, the prevalence of diarrhea in diabetes has been reported to range from 8% to 22%, underscoring its clinical relevance (7, 9, 12, 18). However, the underlying mechanisms of diarrhea differ between T1D and T2D, with a higher prevalence and distinct pathophysiology observed in T1D (33). In T1D, diarrhea typically arises in individuals with long-standing disease and is often attributed to autonomic neuropathy, altered intestinal motility, small intestinal bacterial overgrowth, and an increased incidence of celiac disease. In T2D, glucose-lowering medications, particularly metformin, are well-known contributors to diarrhea (5, 10). However, a subset of T2D patients experience chronic or intermittent diarrhea independent of pharmacologic therapy, which is associated with inflammation, altered gut microbiota, and epithelial dysfunction (10). Notably, one study reported that 40% of T2D patients with non-metformin-related diarrhea met criteria for irritable bowel syndrome (IBS)-diarrhea, suggesting a colonic origin (20).

Preclinical models have been instrumental in elucidating the mechanisms underlying diabetic enteropathy. The streptozotocin (STZ)-induced T1D model, which mimics insulin deficiency through selective β-cell destruction, recapitulates key features of human T1D, including intestinal fluid accumulation and impaired sodium absorption (11). In contrast, STZ-based T2D models commonly combine a high-fat diet (HFD) to induce insulin resistance with subsequent low-dose STZ administration to induce partial β-cell loss. This approach yields a non-autoimmune diabetic phenotype characterized by elevated fasting glucose, glucose intolerance, and moderate hyperinsulinemia, reflecting the metabolic profile of human T2D (11, 25). While overt diarrhea is uncommon in T2D models, they often exhibit epithelial and inflammatory alterations relevant to T2D pathophysiology, including intestinal barrier dysfunction, and dysregulated expression of epithelial ion transporters, such as sodium and chloride channels (30, 35).

A major contributor to intestinal fluid homeostasis is the Na^+^/H^+^ exchanger 3 (NHE3), which mediates the bulk of sodium uptake at the apical membrane of enterocytes in the small intestine and proximal colon. Loss-of-function studies underscore its importance: NHE3-deficient mice develop chronic diarrhea and *SLC9A3* mutations in humans cause congenital sodium diarrhea (17, 32). Reduced NHE3 activity and membrane retention occur in STZ-induced T1D rodent models (15). In contrast, in T2D, metformin-induced AMPK activation can promote NHE3 phosphorylation and ubiquitination, leading to increased endocytosis and reduced surface expression (13). However, whether total NHE3 protein levels are altered in these settings, and whether such changes differ between T1D and T2D, remains unclear, representing a major gap in our understanding of diabetic diarrhea.

We recently demonstrated that human enteroids derived from T1D patients exhibit reduced NHE3 expression and enhanced agonist-stimulated fluid secretion, further supporting the hypothesis that impaired sodium absorption contributes to diarrhea in diabetic enteropathy (3). However, the mechanisms governing NHE3 expression and stability in diabetes, whether transcriptional, post-translational, or segment-specific, remain poorly understood. In the current study, we investigated the regulation of NHE3 in both STZ-induced T1D and T2D mouse models, with a focus on the intestinal segments affected and molecular pathways affecting protein stability. We identified the endosomal adaptor protein SNX17, known for binding FERM-domain cargo motifs and mediating membrane protein recycling, as a novel regulator of NHE3, implicating it in the pathogenesis of T1D-associated small intestinal fluid secretion (26). Through combined in vivo and in vitro analyses, we determined the mechanisms by which NHE3 expression is controlled in the diabetic intestine and showed that distinct mechanisms operate in T1D versus T2D.

## Materials and Methods

Reagents were obtained from Sigma-Aldrich (St. Louis, MO) unless otherwise stated. Glutathione-Sepharose 4B resin (#17-0756-01) was from GE Healthcare. NHS-SS-biotin (#PG82077) was from Pierce Chem. 2,7-Bis(2-carboxyethyl)-5-carboxyfluorescein acetoxymethyl ester (BCECF-AM) (#B1170), SNARF-4F(5-(and-6)-carboxylic acid acetoxymethyl ester) (#S-23920), nigericin (#N1495), and Hoechst 33342 (#62249) were from Invitrogen (Carlsbad, CA). Mouse anti-HA (#MMS-101R) was from Covance, Inc. (Princeton, NJ). Mouse anti-GST (#MA4-004) was from Cell Signaling Technology, Inc. (Danvers, MA). Rabbit polyclonal antibodies against NHE3 were either custom-made (IF and tissue lysate analyses) as previously described (16, 19) or obtained from Novus Biologicals (#NBP1-82574; 1:250) for use in cell-based assays. Rabbit anti-SNX17 antibodies (#NBP1-92417, 1:100) and mouse polyclonal anti-SNX17 antibodies used for PLA (#H0009784-B01P, 1:100) were also purchased from Novus Biologicals. IRdye-700- or IRdye-800-conjugated goat anti-mouse or goat anti-rabbit secondary antibodies were from Rockland Immunochemicals Inc. (Gilbertsville, PA) and were used with LI-COR (Lincoln, NE) Odyssey system for Western blot analysis. Alexa fluor-488 or −568 conjugated goat anti-mouse or anti-rabbit secondary antibodies were from Invitrogen.

### Animals

Type-1 diabetes (T1D) was induced by a single intraperitoneal (i.p.) injection of streptozotocin (STZ) (185mg/kg, Sigma-Aldrich) to C57BL/6J mice (Charles River Laboratories, Inc, Wilmington, MA). After a six-hour fast, adult (8-12 weeks old) male mice received an i.p. injection of STZ dissolved in a 50mM citrate buffer (citric acid and sodium citrate, pH 4.0) or citrate buffer alone for control mice. These mice were maintained on normal chow diet (NCD) (7913 irradiated NIH-31 modified 6% mouse/rat diet – 15 kg, Envigo, Indianapolis, IN). Blood glucose was checked 7 days later with a glucometer via tail vein using a glucometer (OneTouch Ultra 2 meter) and mice with blood glucose levels ≥ 300mg/dl were considered diabetic. STZ-treated mice that had blood glucose levels < 300mg/dl received a second i.p. injection of STZ after a six-hour fast, and blood glucose was checked again 7 days after the repeat injection. If blood glucose levels at the repeat check were≥ 300mg/dl, the mice were considered diabetic; otherwise, they were excluded from the study (18). To induce a model of type-2 diabetes (T2D), male mice (C57BL/6J) aged 5 – 6 weeks were maintained on a high-fat diet (HFD) (Rodent diet with 60 kcal% fat – D12492, Research Diets, New Brunswick, NJ). Five weeks following the initiation of HFD, the mice received daily intraperitoneal (i.p.) injections of low-dose streptozocin (STZ) (40 mg/kg/day, Sigma-Aldrich) for three consecutive days after a six-hour fast on each day (18, 28). Blood glucose levels were measured using the same method described above. STZ was dissolved in a citrate buffer (citric acid and sodium citrate, pH 4.0). For control mice, age-matched littermates were maintained on NCD (7913 irradiated NIH-31 modified 6% mouse/rat diet – 15 kg, Envigo, Indianapolis, IN) and received daily i.p. injections of citrate buffer for three consecutive days at a similar age as the HFD mice. Blood glucose was checked via tail vein two weeks after STZ or citrate buffer injection.

### Cell culture and construction of SNX17-knockdown cells

Human intestinal polarized epithelial cell lines Caco-2bbe and SK-CO15 were used for these studies (31, 33, 34). Cells were grown on membranes (Transwells or filterslips; Corning, Corning, NY) and maintained in DMEM supplemented with 10% FBS, penicillin (50 mU/ml), streptomycin (50 μg/ml) (22). For SK-CO15 cells, the medium was further supplemented with 1 mM sodium pyruvate, 15 mM HEPES, and 1× nonessential amino acids. Stable SNX17 knockdown (SNX17-KD) cells (SK-CO15 and Caco-2bbe) were generated using lenti-shRNA constructs. In brief, gene sequence–specific shRNA clones were constructed within the lentivirus plasmid vector pLKO.1-puromycin (Sigma-Aldrich). Two shRNA constructs were tested to generate lentiviral transduction particles: TRCN0000065163 (#1) GCCTATAACATTCACGTGAAT; TRCN0000065165 (#2) CCAGGCTATCATGATGAGCAT (Sigma-Aldrich). The production of lentiviral particles, lentiviral transduction, and generation of stable cell lines were performed as described previously (23). Knockdown of protein expression was verified by Western blot analysis. As a control, cells were transduced with a lentivirus plasmid vector containing shRNA that does not match any known human gene (Sigma-Aldrich). Infected cells were maintained under the selection pressure of puromycin.

### NHE3 activity measurement in intact mouse jejunum by Two-Photon Microscopy

Na^+^/H^+^ exchange activity in intact mouse jejunum was determined as dimethyl amiloride-sensitive rates of Na^+^-dependent alkalinization using a 2-photon microscope (MRC-1024MP; Bio-Rad, Hercules, CA) and the pH-sensitive dye, SNARF-4F acetoxymethyl ester, as we have described previously (19). Full-thickness jejunum was studied starting 1 cm distal to the ligament of Treitz.

### Measurement of NHE3 activity in cell lines

SK-CO15 and Caco-2 cells grown on Transwell cell culture inserts were used to measure NHE3 activity using the intracellular pH-sensitive dye 2′,7′-Bis(2-carboxyethyl)-5(6)-carboxy fluorescein acetoxymethyl ester (Invitrogen, Carlsbad, CA) and a computerized fluorometer, as described (23, 24). Na^+^/H^+^ exchange rates (H^+^ efflux) were calculated as the initial rates of Na^+^-dependent change in pHi over approximately 1 minute (ΔpH/Δmin). HOE-694 (50 µM) was included in TMA and Na solutions to inhibit potential contributions of NHE1, NHE2, and NHE8 to the NHE activity measured.

### Proximity Ligation Assays

In situ PLA was carried out with the Duolink Detection Kit (#92008, Bethyl Laboratories). Briefly, seven days post confluence, SK-CO15 cells were fixed in 4% PFA for 30 mins, followed by blocking as per the manufacturer’s instructions. The cells were incubated with rabbit NHE3 and mouse SNX17 primary antibodies overnight at 4°C, followed by incubation with secondary antibodies conjugated with the PLA probe at 37°C for 1 h as recommended by the manufacturers. Then, ligation and amplification were performed (Duolink detection kit Orange, 555 nm). Finally, a mounting medium with 4,6-diamidino-2-phenylindole (DAPI) was used. Four non-overlapping fields of view per well were identified, and photomicrographs were acquired under each experimental condition. The images were acquired using ×40 oil immersion objective on an FV3000 confocal microscope (Olympus, Tokyo, Japan) with Olympus FV31S-SW software and processed with Fiji (ImageJ-2020) (NIH). To discriminate PLA puncta from the background fluorescence, the same manually selected threshold was applied to all images. The number of nuclei (DAPI+; ∼100 per field of view per condition) and total puncta (white spots) were counted using the Duolink1 Image Tool Software (Olink Bioscience) from each of the four fields of view and averaged for each experimental condition for statistical comparison. A total of five biological replicates were performed. One-way ANOVA followed *by a priori* comparisons with Tukey’s test or Student’s t-tests were conducted, as appropriate.

### RNA isolation and Quantitative Real-Time PCR

Total RNA was extracted from mouse mucosal samples using Qiagen RNeasy kits according to the manufacturer’s protocol. Complementary DNA was synthesized from 1 to 2 μg of RNA using SuperScript VILO Master Mix (Life Technologies). Quantitative real-time PCR was performed using Power SYBR Green Master Mix (Life Technologies) on a QuantStudio 12K Flex real-time PCR system (Applied Biosystems, Foster City, CA). Each sample was analyzed in triplicate, and 5 ng RNA-equivalent complementary DNA was used for each reaction. Commercially available primer pairs from OriGene Technologies (Rockville, MD) were used: **Mouse:** *Nhe3*: MP215958; *Snx17:* MR207515; and glyceraldehyde-3-phosphate dehydrogenase (*Gapdh*): MP205604**; Human:** *NHE3*: HP207529; *SNX17:* HP211121; and *GAPDH*: HP205798. The relative fold changes in mRNA levels of genes were determined by using the 2 ^-ΔΔCT^ method with human GAPDH, RNA simultaneously studied and used as the internal control for normalization, and results are shown as fold change compared to control. Statistical comparisons were performed on raw ΔCt values to preserve group variance before normalization.

### Recombinant protein purification

Mouse SNX17 cDNA was kindly provided by Ralph Böttcher (Max Planck Institute of Biochemistry, Germany). GST–SNX17-Full length (FL) was constructed and purified as previously described(23). cDNAs encoding full-length SNX17 were generated by PCR and inserted into pGEX-4T-1 (GE Healthcare) for expression of GST-fused recombinant proteins. The constructs were transformed into the BL21(DE3) strain (EMD Millipore). When the bacterial culture reached an optical density (OD) of ∼0.8, protein expression was induced with 0.3 mM isopropyl-β-D-thiogalactoside at 16°C overnight. GST-tagged protein was purified in a gravity-flow column following the instructions from GE Healthcare. Purified proteins were concentrated with Amicon Ultra-15 Centrifugal filter units (EMD Millipore) supplemented with 10% glycerol and 10 mM DTT, and protein concentrations were measured via Bradford assays.

### Immunoprecipitation (GSH resin pulldown)

Immunoprecipitation experiments were performed using lysates from control or HA-NHE3 infected Caco-2bbe cells as previously described (23). Briefly, 1 nmol of recombinant GST-tagged protein was used as bait. As prey, 1 mg of cell lysate from HA-NHE3 or HA-NHE3-F760A (A^760^) mutant generated using the QuikChange site-directed mutagenesis kit (Agilent Technologies) was used. The volume of the final mixture was adjusted to 500 μl with the lysis buffer (25 mM HEPES, pH 7.4, 150 mM NaCl, 50 mM NaF, 1 mM Na_3_VO_4_, 0.5% Triton X-100 and protease inhibitors). GSH resin (Glutathione–Sepharose 4B resin) was washed with lysis buffer three times. Each bait–prey mixture was mixed with 10 μl of resin and incubated at 4°C for 4 h on a rotating shaker. Resin was washed with the same lysis buffer four times and then eluted with lysis buffer supplemented with 10 mM glutathione. The input and elution samples were analyzed by SDS-PAGE and western blotting.

### Cell Surface biotinylation

Seven days postconfluent SK-CO15 cells were grown on 10-cm-diameter filters (Corning). The cells were then serum-starved for ∼3 h. All subsequent manipulations were performed at 4°C. For surface labeling of NHE3, cells were incubated with 1.5 mg/ml NHS-SS-biotin (biotinylation solution; Pierce Chemical, Rockford, IL) for 30 min and repeated once, solubilized with lysis buffer, and then incubated for 6 h with streptavidin-agarose beads. Western analysis and the quantification of the surface fraction and surface/total ratio were performed as described previously with normalization to GAPDH (1, 23).

### Total NHE3 degradation assay

Postconfluent control and SNX17-KD SK-CO15 and Caco-2bbe cells were grown on Transwell filters and treated with 100 μM cycloheximide for the indicated time points. Cells were lysed in PBS with 1% (vol/vol) Triton X-100, and NHE3 levels were determined by quantitative Western blotting. β-Actin fluorescence intensity was used to normalize the detected levels of NHE3. The untreated controls were set to 100%, and the level of detected NHE3 was calculated as the percentage of untreated control for each time point.

### Statistics

Statistical significance was assessed by one-way ANOVA with Tukey’s test or Student’s t-tests. Results are presented as mean ± SEM. A *P-*value of less than or equal to 0.05 was considered significant.

## Results

### Increased luminal fluid content in the small intestine of T1D and T2D mice and reduced jejunal NHE3 activity in T1D but not in T2D mice

The occurrence of intermittent diarrhea is common in patients with diabetes, and reduced fluid absorption is established in the STZ-T1D mouse model (15). We used STZ-induced type 1 diabetic (T1D) and high-fat diet-STZ type 2 diabetic mice to investigate the mechanism contributing to diabetic diarrhea. A blood glucose level of >300 mg/dL was the criterion used for successful induction of diabetes in all STZ groups (Figure 1A). The STZ-treated mice had dilated small intestines compared to the control mice. To assess the differences in luminal fluid content between the control and STZ mice, we calculated the ratio of wet (water) weight to dry weight of the entire small intestine. The small intestine wet/dry weight ratio was significantly increased in STZ-treated mice compared to controls, with no significant difference between T1D and T2D groups (Figure 1B). Fluid absorption as well as Na+/H+ exchanger-3 (NHE3) activity is known to be reduced in the small intestine of the STZ T1D mouse model(15). To further understand the fluid defects in STZ mice, we examined multiple aspects of NHE3. The Na+/H+ exchange activity was measured in intact mouse jejunum by 2-photon microscopy using the pH-sensitive dual-emission dye, SNARF-4F. This showed reduced jejunal NHE3 activity in T1D (ΔpH/min: 0.25 ± 0.03 vs 0.13 ± 0.01 control and T1D, respectively, *P* < 0.05) (Figure 1*C*). In contrast, the effects on NHE3 were different in T2D in which there was no significant change in jejunal NHE3 activity (ΔpH/min: 0.25 ± 0.03 control vs 0.26± 0.05 T2D; *P*=NS) (Figure 1*C*).

**Figure 1.**
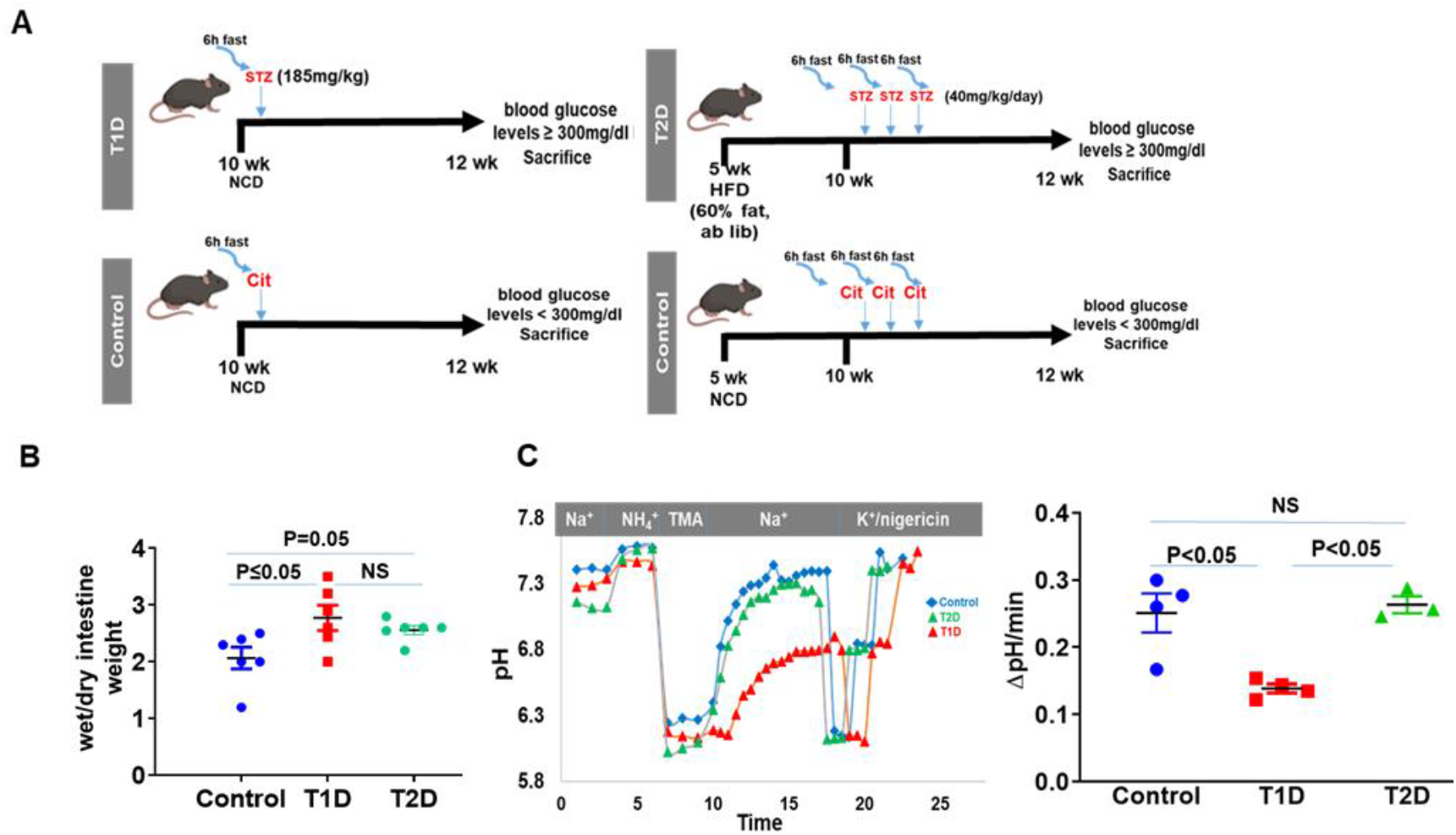
Increased intestinal fluid content and decreased NHE3 activity in the small intestine of STZ T1D mice. *A:* Study design for diabetes induction in male C57BL/6J mice involved two models. For T1D, a single i.p. injection of streptozotocin (STZ) (185 mg/kg) was given to induce diabetes, with control mice receiving citrate buffer. Blood glucose was measured after 7 days, with levels ≥ 300 mg/dl indicating diabetes. For T2D, mice were placed on a high-fat diet (HFD) starting at 5-weeks of age. At 10^th^ weeks they received daily i.p. injections of low-dose STZ (40 mg/kg) for three days after fasting. Control mice received citrate buffer under the same conditions. Blood glucose was checked two weeks post-injection. *B:* Ratio of wet/dry weight of entire small intestine showing increased luminal fluid in STZ treated mice compared to control. Results are shown as the means ± SEM; n=6 animals per group. NS, not significant. *C:* Murine jejunal pHi determination over time is superimposed on the experimental protocol. The tissue was loaded with 20 μM SNARF-4F, and EIPA-sensitive basal NHE3 activity in jejunum was determined by two-photon microscopy as the initial rate of Na^+^-induced alkalinization after an acute acid load. Fifty micromolar HOE-694 was present to eliminate the contribution of NHE1 and NHE2 activities to initial rates. Dot plot (right) showing decreased basal NHE3 activity in T1D mouse jejunum compared to control and T2D group. Data points in *(B)* and *(C)* represent number of animals studies. Results are shown as the means ± SEM; n=3-6 animals per group. NS, not significant. Comparison done with ANOVA.

### Reduced NHE3 expression in the duodenum and jejunum of STZ-induced T1D mice is not associated with changes in mRNA expression, whereas in T2D mice, there is reduced NHE3 mRNA expression in the proximal colon

To assess the mechanism of decreased NHE3 activity in T1D, we analyzed NHE3 expression across the intestine of STZ mice along with corresponding mRNA levels. In T1D mice, a significant reduction in NHE3 protein levels was observed in the duodenum and jejunum, as demonstrated by both western blot analysis of mucosal lysates and immunofluorescence (IF) staining of tissue sections (Figures 2A, B). In the ileum, IF staining (mid ileal region) revealed a decrease in membrane-localized NHE3 with a concurrent increase in intracellular pools. However, the corresponding reduction in total NHE3 protein levels did not reach statistical significance by western blot (Figures 2A–C). There was no change in proximal colonic NHE3 protein expression in T1D mice (Figure 2B). In addition, there was no change in duodenal, jejunal, and proximal colonic NHE3 mRNA in T1D mice, although NHE3 mRNA was reduced in the ileum of these mice (Figure 2C). In contrast, T2D mice displayed a distinct pattern of NHE3 regulation, with no significant changes in NHE3 expression in the small intestine, while there was a marked reduction in both NHE3 mRNA and protein expression in the proximal colon. This reduction was consistently demonstrated by histological staining and western blot analysis, indicating that reduced NHE3 transcription contributes to reduced NHE3 protein expression in T2D mice, specifically in the colon (Figures 2A-C). Collectively, these findings suggest that in T1D mice, the reduction in NHE3 expression within the duodenum and jejunum is primarily due to post-transcriptional mechanisms or impaired protein stability, as mRNA levels remain unchanged. In the ileum of T1D mice, transcriptional downregulation also occurs; however, its impact on overall NHE3 protein expression appears minimal. In contrast, the small intestine of T2D mice shows no significant change in NHE3 expression, whereas a marked reduction was observed in the proximal colon. This colonic decrease in NHE3 protein in T2D mice is driven by transcriptional downregulation.

**Figure 2.**
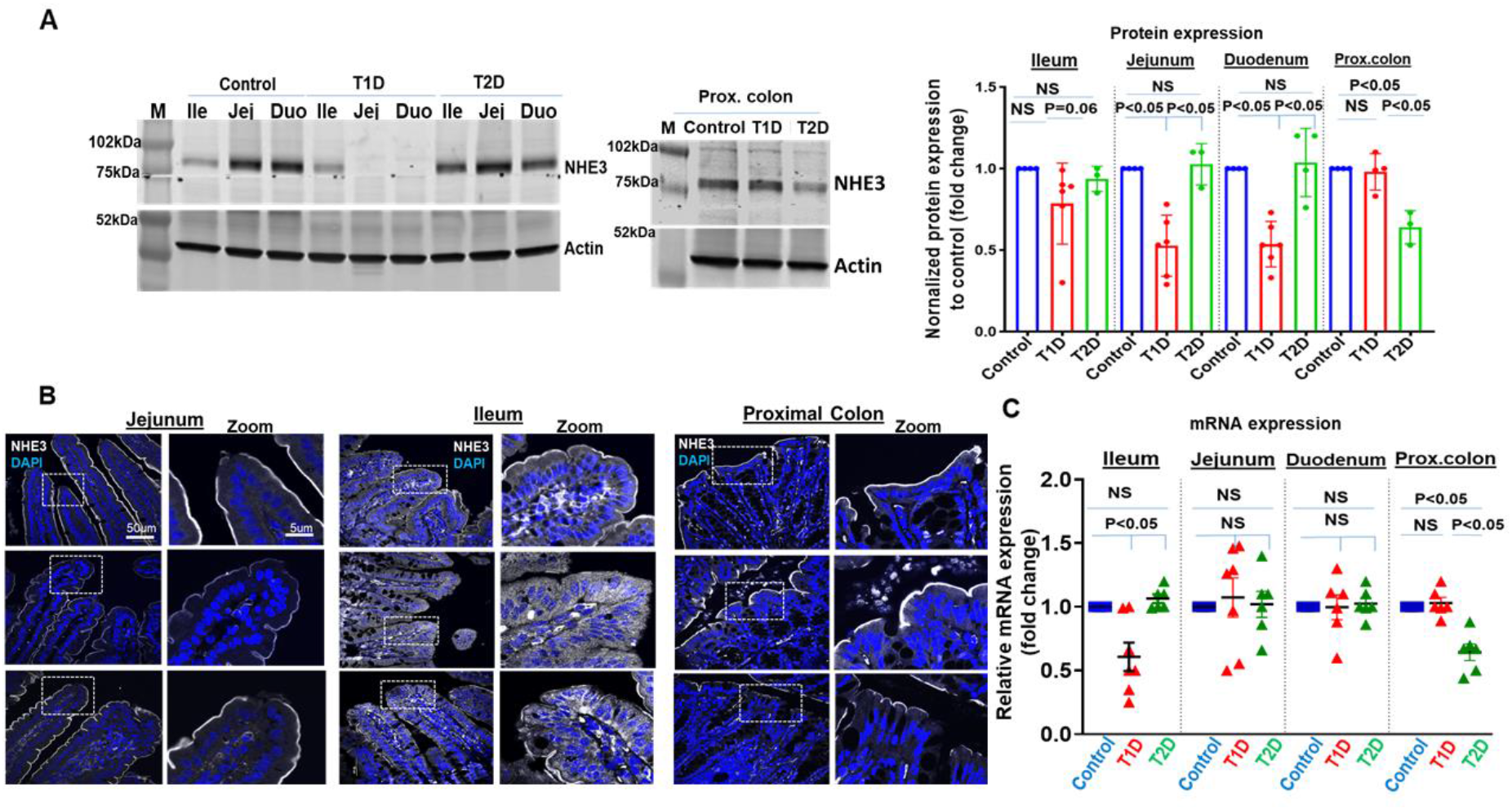
Total NHE3 expression decreases differentially in T1D and T2D. *A:* A representative western blot demonstrates the expression levels of NHE3 in lysates prepared from the duodenum, jejunum, ileum, and proximal colon of control and STZ-treated mice. Quantitative analysis (shown to the right) shows a significant decrease in NHE3 protein expression in the duodenum and jejunum of T1D mice when compared to control and T2D groups. In contrast, in the proximal colon, a reduction in NHE3 expression is observed in T2D mice compared to both T1D and control mice. Results are means ± SEM; n=3-6 animals per group were analyzed. NS, not significant. *B:* Representative confocal image showing reduced NHE3 expression (white) in the jejunum and ileum of T1D mice and in the proximal colon of T2D mice, compared to controls. Nuclei are stained blue. Confocal was performed on 3-6 animals per group, and representative images from each group are shown. *C:* Relative *Nhe3* mRNA expression in the duodenum, jejunum, ileum, and proximal colon of control and STZ-treated mice. Data are normalized to control (set to 1.0) and presented as fold change (mean ± SEM; n = 3-6 mice per group). Statistical comparisons were performed using unpaired t-tests on ΔCt values; NS indicates not significant. Comparisons in *A* and *C* are done with ANOVA.

### Decreased expression of NHE3 in the small intestine of STZ T1D mice is due to reduced endosomal SNX17 expression

Given that the inhibition of NHE3 activity in the small intestine of T1D mice was associated with reduced total protein expression, we next sought to investigate the molecular mechanisms contributing to this decrease. NHE3 activity is highly regulated by intracellular trafficking involving the endosomal proteins. We previously reported that the <u>P</u>ostsynaptic density protein (PSD95), <u>D</u>rosophila disc large tumor suppressor (Dlg1), and <u>Z</u>onula occludens-1 protein (ZO-1) (PDZ) domain-containing endosomal scaffolding protein SNX27 and its interaction with the PDZ-binding motif of NHE3 are important for plasma membrane localization and recycling of NHE3 in intestinal epithelial cells. However, this interaction does not regulate the total cellular expression of NHE3 (23). In contrast, SNX17 is another endosomal scaffolding protein, which has been implicated in the trafficking and stability of several membrane proteins (4, 26). We therefore explored the potential involvement of SNX17 in contributing to changes in NHE3 in T1D. As shown in Figures 3A and 3B, both SNX17 mRNA and protein levels were significantly reduced in the duodenum and jejunum of STZ-induced T1D mice compared to controls. In the ileum, SNX17 protein levels were modestly reduced, although there was no significant change in *Snx17* mRNA expression. In contrast, the proximal colon of T1D mice showed no change in SNX17 mRNA or protein levels. Similarly, T2D mice exhibited no significant alterations in SNX17 expression in either the small intestine or proximal colon. These findings suggested that the reduction of SNX17 in the duodenum and jejunum of T1D mice might contribute to the reduced NHE3 protein expression in these segments and warranted further studies of the potential mechanism involved.

**Figure 3.**
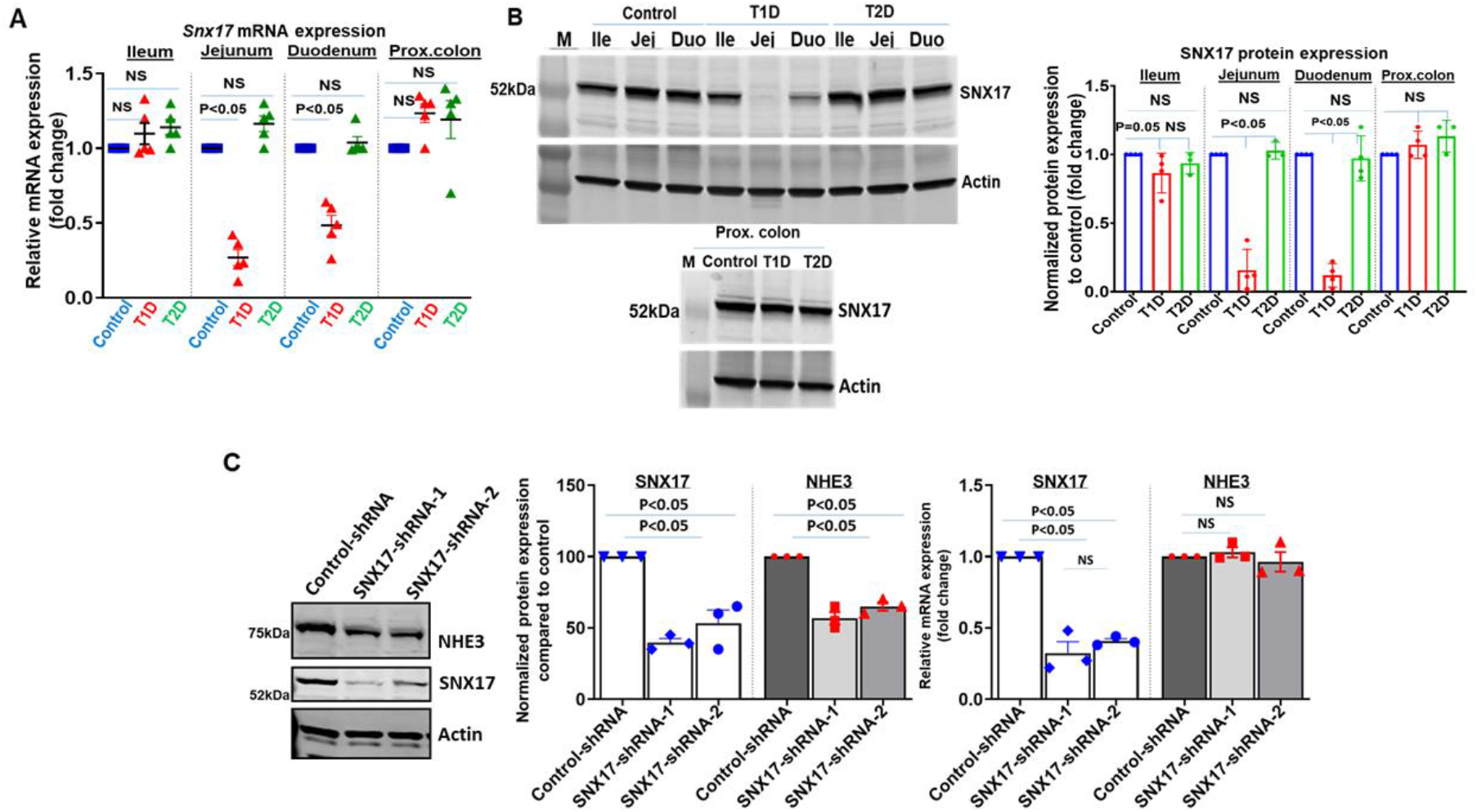
Decreased expression of NHE3 in the small intestine of STZ T1D mice is due to reduced endosomal SNX17 expression. *A:* Relative mRNA expression analysis shows decrease in *Snx17* mRNA expression in duodenum and jejunum but not in ileum of T1D compared to control and T2D mice. Results are means ± SEM; n=3-4 animals per group were analyzed. NS, not significant. *B*: Western blot demonstrating expression of SNX17 in lysates prepared from duodenum, jejunum, ileum, and proximal colon of control and STZ-treated mice. Analysis of multiple western blots showing decrease in SNX17 in duodenum and jejunum of T1D mice compared to control and T2D. Results are means ± SEM; n=3-4 animals per group were analyzed. NS, not significant. *C:* A representative western blot (left) and quantification from 3 independent analyses (middle) shows the effect of two distinct SNX17-targeting shRNAs and a nontargeting control shRNA on SNX17 and NHE3 protein levels in polarized SK-CO15 cells. Stable expression of shRNA was maintained by growing cells in puromycin-containing media. Both SNX17 shRNAs similarly decreased SNX17 and NHE3 levels compared with control shRNA. Actin served as an internal control. Results are means ± SEM. Corresponding qPCR data (right) show the impact of the same shRNAs on *SNX17* and *NHE3* mRNA expression. Results are means ± SEM; n=3 independent analyses. NS, not significant. Comparisons done with ANOVA.

To determine whether SNX17 influences NHE3 protein expression, we generated a stable SNX17 knockdown (SNX17-KD) in the human colon epithelial cell line SK-CO15, which endogenously expresses NHE3.

Knockdown was achieved using a lentiviral delivery system encoding short hairpin RNA (shRNA) targeting SNX17 (Figure 3C). Two independent shRNA constructs were tested, each producing a substantial reduction in SNX17 protein levels, approximately 65% ± 4 (shRNA-1) and 55% ± 5 (shRNA-2), compared to control shRNA-expressing cells. Knockdown efficiency was maintained under puromycin selection. Importantly, SNX17 depletion led to a significant reduction in NHE3 protein levels, with shRNA-1 and shRNA-2 resulting in approximately 45% ± 4 and 35% ± 6 decreases, respectively, relative to control cells (Figure 3C). Quantitative PCR confirmed that *NHE3* mRNA levels remained unchanged in SNX17-KD cells (Figure 3C, right), indicating that the downregulation of NHE3 occurred post-transcriptionally and was likely due to impaired protein trafficking or stability. These in vitro results pointed to a possible role for SNX17 in maintaining NHE3 protein expression, prompting us to investigate whether alterations in SNX17 expression might underlie the NHE3 deficiency observed in T1D mice.

### SNX17 interacts with and is necessary for basal NHE3 activity

Interaction between SNX17 and NHE3 was assessed using an in situ proximity ligation assay (PLA), which generates an enhanced fluorescent signal when two proteins physically associate (2, 8). The PLA was performed on SK-CO15 cells grown as a polarized model on Transwell inserts. As shown in Figure 4A, NHE3-SNX17 interaction occurred under basal conditions, as evidenced by the enhanced signal (white dots) compared to the monolayer used as a negative control. To determine that the reduced SNX17 expression affects NHE3 basal activity, we also measured NHE3 activity in SNX17-KD SK-CO15 cells. Consistent with reduced expression of NHE3 in SNX17-KD cells, NHE3 basal activity was significantly reduced in SNX17-shRNA (shRNA-1: 0.12 ΔpH/min ±0.02; shRNA-2: 0.16 ΔpH/min ±0.07) expressing cells compared to control shRNA (0.23 ΔpH/min ±0.04) (Figure 4B). Together, these results demonstrate that SNX17 physically associates with NHE3 under basal conditions and is necessary for sustaining its basal transport activity.

**Figure 4.**
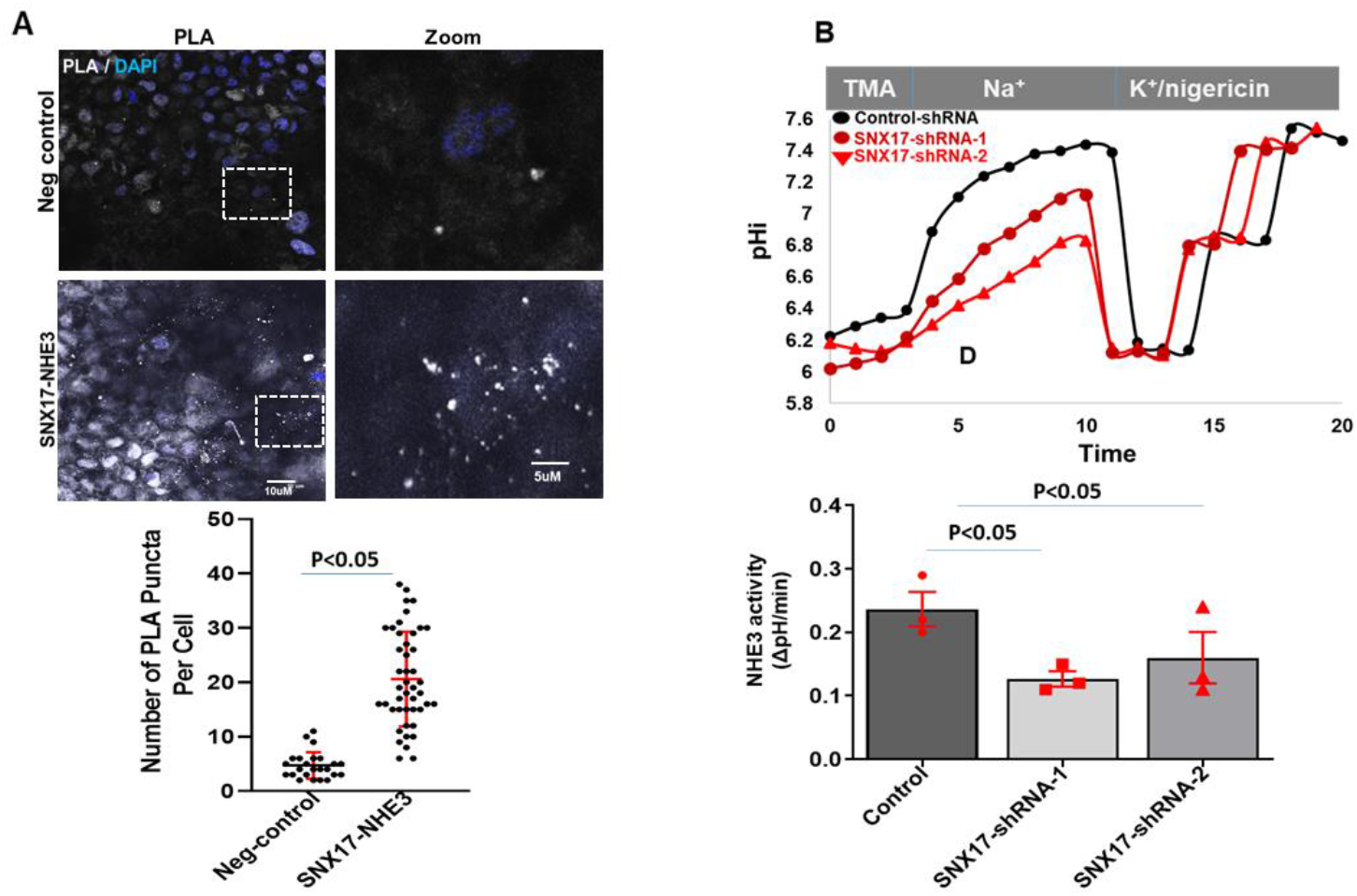
SNX17 interacts with NHE3 and regulates its basal NHE3 activity. ***A:*** A representative image of in situ proximity ligation assay (PLA) and quantitation (below) demonstrates the interaction between NHE3 and SNX17. Polarized SK-CO15 monolayers were utilized to determine the interaction between endogenously expressing NHE3 and SNX17 using specific antibodies for both proteins. Interaction was indicated by white dots, while nuclei were stained with DAPI (blue). The number of white puncta (mean ± SEM) was quantified from over 100 representative cells across multiple monolayers. The experiment was independently repeated three times (n = 3), and the combined results are presented here. Monolayers that lacked primary antibodies served as negative controls for background PLA signal. Student’s paired T tests. *B:* Basal NHE3 transport activity was measured as the initial rates of Na^+^-dependent intracellular alkalinization in control and SNX17shRNA-expressing monolayers using the pH sensitive dye BCECF. Fifty micromolar HOE-694 was maintained throughout to eliminate the contributions of NHE1 and NHE2 activities to initial rates. A single experiment is shown above, and below are shown means±SEM., n=3 independent experiments were analyzed. Comparison done with ANOVA.

### SNX17 regulates NHE3 protein stability but not its percent surface distribution

To determine whether SNX17 influences NHE3 surface expression as the mechanism for the reduced NHE3 activity when SNX17 is KD, we investigated the apical abundance of NHE3, as a percentage of total NHE3 expression, in polarized SKCO-15 epithelial cells using surface biotinylation. Although we previously showed that SNX27 modulates NHE3 localization at the apical membrane under basal conditions, the role of SNX17 in NHE3 apical membrane trafficking is not known. SK-CO15 cells stably expressing either control or SNX17-shRNA were grown as apically outward-facing polarized monolayers on Transwell filters and subjected to apical biotin labeling to quantify membrane-localized NHE3. As shown in Figure 5A, SNX17-KD resulted in a marked decrease in the absolute amount of NHE3 at the apical membrane compared to control cells. However, when normalized to total NHE3 levels, the ratio of surface-to-total NHE3 remained unchanged between SNX17-KD and control cells (control-shRNA: 20% ± 0.04; SNX17-shRNA-1: 19% ± 0.05; P = NS, n=8). This indicates that the SNX17-related decrease in surface NHE3 reflects the overall reduction in total NHE3 protein levels rather than a change in trafficking. To further investigate the mechanism underlying the reduced NHE3 protein levels in SNX17-deficient cells, we assessed whether SNX17 regulates the stability or degradation rate of NHE3 using SK-CO15 cells. A cycloheximide chase assay was used to inhibit new protein synthesis, and total NHE3 expression levels were measured at time points between baseline and 48 hours by western blot analysis. As shown in Figure 5B, in control-shRNA cells, NHE3 exhibited a half-life of approximately 16 hours, consistent with a relatively stable membrane protein as reported previously in PS120 fibroblasts (6). In contrast, SNX17-KD cells displayed a significantly accelerated degradation of NHE3, with an estimated half-life of ∼8 hours (Figure 5B). These results demonstrate that SNX17 does not directly regulate the trafficking of NHE3 to the apical membrane or alter its percent surface expression under basal conditions. Instead, SNX17 is necessary to establish the overall half-life of NHE3.

**Figure 5.**
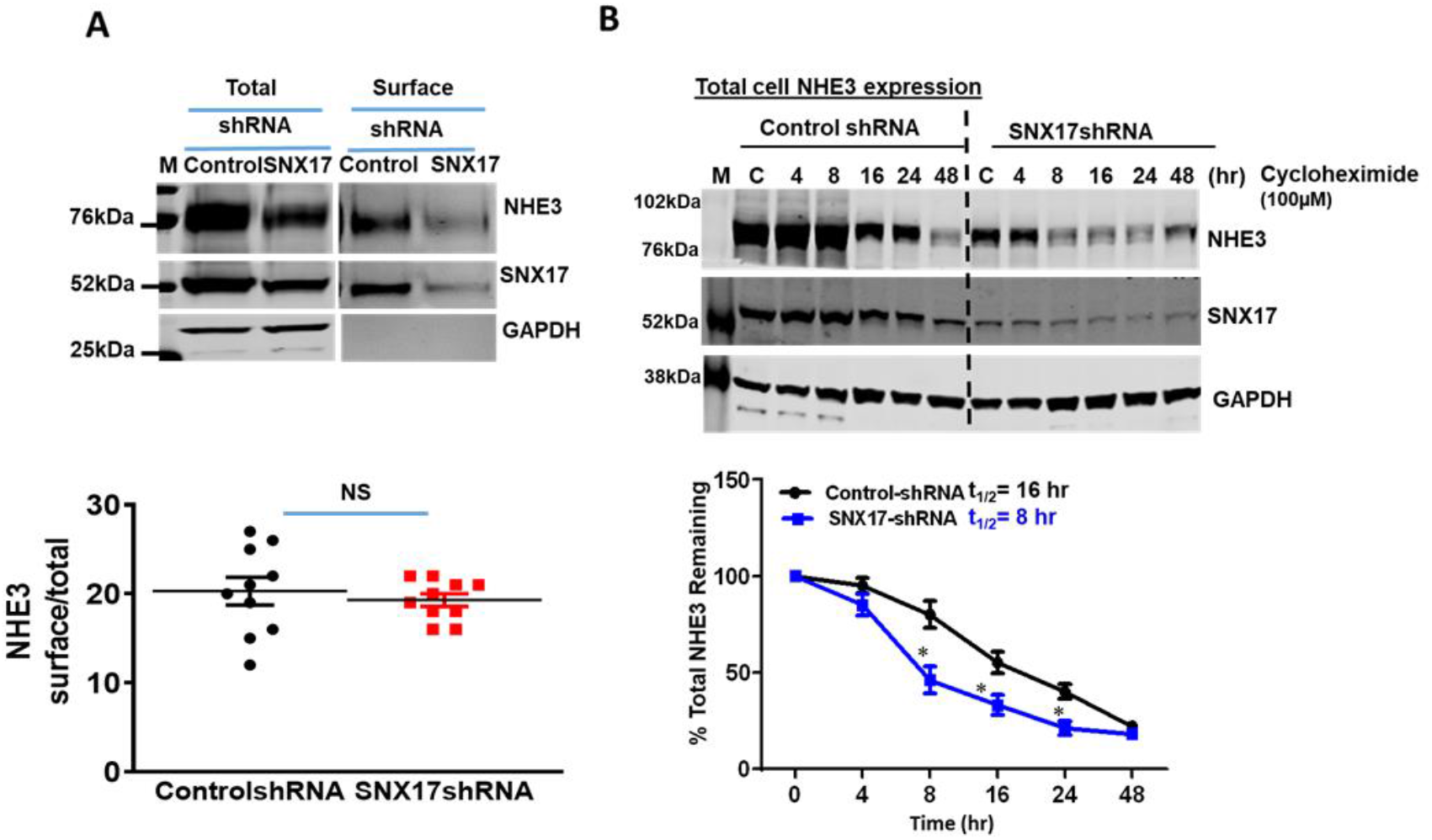
NHE3 half-life, but not percent surface expression, is regulated by SNX17. *A:* Representative western blot and quantification (below) of multiple analyses show no change in the surface/total ratio of NHE3 in SNX17 shRNA expressing cells compared with the control. Cell surface biotinylation studies were performed on monolayers from polarized SK-CO15 cells expressing control and SNX17 shRNA. Results are shown as the means ± SEM; n=8-9 experiments. NS, not significant. *B:* Stability of total NHE3 in control and SNX17 shRNA expressing SK-CO15 cells measured in presence of cycloheximide (100µM) to prevent new protein synthesis. A representative blot and quantitation (below) show shorter half-life of NHE3 in SNX17shRNA-expressing cells compared to control. Densitometric analyses of multiple western blots were performed to calculate the percentage of NHE3 remaining over indicated periods. Results are presented as mean ± SEM from 3-4 independent experiments. *P < 0.05 versus control at the same time point. Comparison done with Student’s paired T tests (*A*) and with ANOVA (*B*).

### SNX17 interacts with the NHE3 C-terminus at a conserved NPxY motif

Sorting nexin 17 is an adaptor molecule that contains a C-terminal noncanonical FERM-like domain (4.1 protein, Ezrin, Radixin, and Moesin), which recycles cargo by binding to the tyrosine/phenylalanine-based-NPxY/F sorting motif (Figure 6A) that is present within the intracellular cytosolic domain of numerous integral membrane proteins (27). To identify the specific binding site between SNX17 and NHE3 and whether this binding regulates NHE3 activity and expression, we performed NHE3 sequence alignment and identified a conserved <u>NPx**F**</u> motif at 757-760aa in the intracellular C-terminal domain of NHE3 (Figure 6A). This site of NHE3 (HA-NHE3 (rabbit)) was mutated, **F**760**A** (NPx**A**^760^). The HA-NHE3-wt or HA-NHE3-A^760^ were transiently transfected into Caco-2bbe cells. We first examined whether the NHE3-A^760^ mutant affected NHE3 binding to SNX17. This was achieved by generating GST fusion proteins of the wild-type-SNX17. Purified GST-SNX17-full length was used to pull down NHE3 from lysates of Caco2-bbe cells transiently expressing HA-NHE3-wt or HA-NHE3-A^760^. GST-SNX17 successfully pulled down HA-NHE3-wt, but not HA-NHE3-A^760^ (Figure 6B). This suggests that SNX17 interacts with NHE3 and involves the conserved NPxF^760^ consensus sequence. Next, we assessed the effect of this mutation on NHE3 basal activity using Caco-2bbe cells transiently expressing (adenovirus) the NHE3 constructs. The SNX17 binding deficient mutant of NHE3 showed a significantly lower basal activity as compared to wt-NHE3, highlighting the importance of SNX17 interaction in maintaining NHE3 basal activity (NHE3-wt: ΔpH/min: 0.28±0.11, NHE3-760A: ΔpH/min: 0.14 ± 0.09, *P* < 0.05) (Figure 6C). Finally, we investigated whether the NHE3-A^760^ mutant affected the overall half-life of NHE3, similar to what was observed in SNX17-KD cells. The half-lives of HA-NHE3-wt and HA-NHE3-A^760^ were determined in Caco-2bbe cells treated with cycloheximide as above. As shown in Figure 6C, the HA-NHE3-A^760^ had a half-life of approximately 8h, significantly shorter than the 16h half-life observed for HA-NHE3-wt. Overall, these results suggested that binding of SNX17 to NHE3 at the conserved 757-760aa NPx**F** site is necessary for maintaining the NHE3 half-life and setting its basal activity.

**Figure 6.**
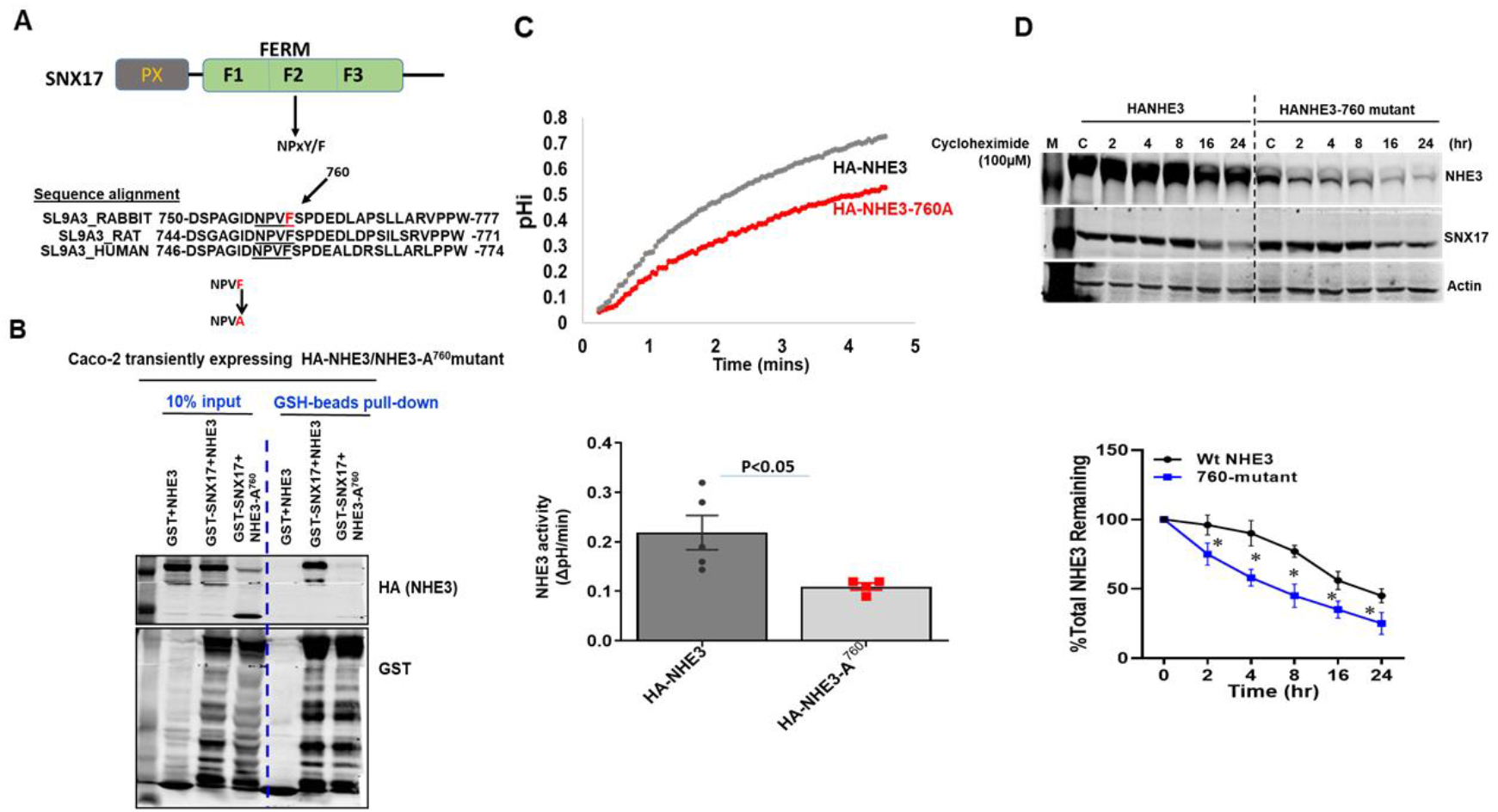
SNX17 interacts with the NHE3 C-terminus at a conserved NPxY motif. *A:* A schematic diagram illustrates the organization of the SNX17 domains, highlighting both the PX domain and the FREM domains. A sequence alignment of NHE3 across different species, emphasizing the conserved SNX17 binding motif, NPxY/F, located at the C-terminus of NHE3. *B:* GST-pull down assay demonstrating loss of interaction between SNX17 and NPxF-760A mutant of NHE3. GST or GST-SNX17 (full length) were mixed with lysates prepared from Caco-2bbe-HA-NHE3 (wt) or Caco-2bbe-HA-NHE3-A^760^ and then subjected to pull down with GSH resin. Samples were analyzed against HA (NHE3) and GST. A representative blot from four independent pull-downs is shown. *C:* A representative trace and quantitation of multiple analyses show a decrease in NHE3 activity in the NHE3-A^760^ mutant-infected Caco-2bbe cell monolayer when compared to the wild-type NHE3. Fifteen days post confluent Caco-2bbe cells grown on Transwell inserts were transiently infected with either adenovirus HA-NHE3 or HA-NHE3-A^760^. NHE3 activity was assessed by measuring the initial rates of Na^+^-dependent intracellular alkalinization utilizing the pH-sensitive dye BCECF in the presence of HOE-694 (50µM) to exclude the contributions of NHE1 and NHE2 activities during the initial rate measurements. Results are means±SEM, n=3-5 independent experiments were analyzed. *D:* Half-life of 760A mutant of NHE3 (NHE3-A^760^) compared to wild-type NHE3 in Caco-2bbe cells. Fifteen days post confluent Caco-2bbe cells were infected with either HA-NHE3 or HA-NHE3-760A mutant and the total NHE3 remaining was determined in the presence of cycloheximide (100µM) as explained above. A representative blot and quantitation (below) show a shorter half-life of HA-NHE3-A^760^ compared to HA-NHE3. Results are means ± SEM; n=3-4 independent experiments were analyzed. *P < 0.05 versus control at the same time point. Comparison done with Student’s paired T tests (*C*) and with ANOVA (*D*).

## DISCUSSION

Diarrhea is a common gastrointestinal complication in individuals with diabetes, though its underlying mechanisms remain incompletely understood. Contributing factors such as autonomic neuropathy, altered motility, gut dysbiosis, and hormonal imbalances are all known to impact intestinal ion transport. Among these, impaired sodium absorption has emerged as a major contributor to fluid imbalance in nearly all diarrheal diseases, including diabetic enteropathy. In this study, we investigated the regulation of NHE3, a major determinant of intestinal sodium and water absorption, in T1D and T2D. Our findings reveal that although both T1D and T2D mice exhibited increased luminal fluid content and reduced NHE3 protein expression, the mechanisms driving NHE3 dysregulation are distinct and segment-specific. Moreover, we identify, for the first time, a novel role for the endosomal scaffolding protein SNX17 in regulating total NHE3 protein stability in the small intestine, specifically in the context of T1D.

Consistent with previous studies, we observed a significant reduction in NHE3 expression and activity in the small intestine of STZ-induced T1D mice (15). However, unlike prior reports, which emphasized mislocalization or defective apical targeting of NHE3 (attributed to altered interactions with PDZ domain-containing scaffolds such as Na^+^–H^+^ exchanger regulatory factor (NHERF)1/2), our data demonstrate a robust decrease in total NHE3 protein expression without a corresponding reduction in NHE3 mRNA. This implicates a post-translational mechanism driving NHE3 loss in T1D. Intriguingly, we found that the endosomal sorting protein SNX17 is markedly downregulated in these same intestinal regions. Functional studies in intestinal epithelial cells confirmed that SNX17 directly binds NHE3, and SNX17-KD reduces NHE3 activity without affecting its apical trafficking. Cycloheximide chase assays further demonstrated accelerated degradation of NHE3 upon SNX17 depletion, identifying SNX17 as a key regulator of NHE3 protein stability.

Although NHE3 has a PDZ domain recognition motif, neither NHERF 1 or 2, nor the endosomal protein SNX27, which also has a PDZ domain, appeared to influence NHE3 stability (23). The current study highlighted the role of another endosomal scaffolding protein, SNX17, in the regulation of NHE3 stability. SNX17 is closely related to the key endosomal trafficking machinery, SNX27-retromer (29). It is a part of a functionally distinct multiprotein complex known as the retriever (21). Unlike SNX27, which primarily handles PDZ domain-containing cargo proteins, SNX17 specifically governs the plasma membrane trafficking of NPxY motif-containing cargos. Mechanistically, SNX17 interacts with the conserved NPxF motif in the cytosolic C-terminus of NHE3. This motif is known to be recognized by the FERM-like domain of SNX17, which governs the recycling and stabilization of membrane proteins such as β1-integrin and members of the LDL receptor family (4, 28). Our findings suggest that NHE3 is regulated by two separate endosomal complexes: SNX27 for apical trafficking and surface expression via PDZ-dependent interactions (23), and SNX17 for maintaining basal protein levels via FERM–NPxF interactions.

Importantly, SNX17-dependent regulation of NHE3 appears to be specific to T1D and not T2D. In T2D mice, SNX17 expression is maintained across all intestinal regions, and NHE3 downregulation is restricted to the proximal colon, where it is accompanied by decreased mRNA expression, suggesting transcriptional repression rather than post-translational degradation. In contrast, T1D is characterized by reduced SNX17 and NHE3 protein expression in the duodenum and jejunum without corresponding changes in NHE3 mRNA, consistent with a post-translational regulatory mechanism. Interestingly, in the ileum of T1D mice, the regulatory profile diverges: SNX17 protein is reduced despite unchanged mRNA levels, NHE3 mRNA is modestly but significantly decreased, yet NHE3 total protein expression, while reduced, is less affected than in more proximal small intestine. This pattern suggests that NHE3 in the ileum may be regulated through dual mechanisms, where transcriptional repression occurs independently of SNX17, and a threshold level of SNX17 loss may be necessary to destabilize NHE3 protein, a threshold not met in this segment. These observations underscore the intestinal segment–specific control of NHE3 and suggest that distinct mechanisms may predominate along different regions of the gut, even within the same disease model. Overall, these results point to disease- and segment-specific regulatory pathways: SNX17-dependent post-translational destabilization of NHE3 in the small intestine during T1D, and transcriptional downregulation of NHE3 in the colon during T2D, potentially driven by inflammation, insulin resistance, or metabolic signaling. Notably, despite preserved NHE3 expression in the small intestine of T2D mice, luminal fluid accumulation was still increased, suggesting additional mechanisms may contribute to fluid loss and warrant further investigation. Similarly, segmental specificity, small intestinal involvement in T1D and proximal colonic changes in T2D, remains poorly understood, but underscores the need for further investigation into region-specific regulation of sodium transporters in diabetic enteropathy. Notably, our models excluded the confounding effects of metformin, a known NHE3 inhibitor (14), allowing us to identify diabetes-intrinsic changes in NHE3 regulation. Clinical studies support the relevance of our T2D findings: even in the absence of metformin, a significant proportion of T2D patients experience chronic diarrhea, with approximately 40% of non-metformin cases meeting criteria for IBS, a condition associated with colonic dysfunction (20). This observation aligns with our identification of proximal colonic NHE3 downregulation in T2D mice, supporting the idea that epithelial transport defects in the colon may contribute to metformin-independent diarrhea in T2D.

Finally, our results are reinforced by human enteroid studies from T1D patients, which demonstrated decreased NHE3 expression and enhanced fluid secretion in the small intestine, highlighting the translational relevance of our findings and further implicating impaired sodium absorption as a central mechanism in diabetic diarrhea (3). Taken together, our data present a novel mechanistic model in which NHE3 dysregulation is disease-type and region-specific: **In T1D**, SNX17 downregulation leads to destabilization and degradation of NHE3 protein in the small intestine, resulting in decreased sodium absorption and fluid accumulation. **In T2D**, NHE3 is transcriptionally repressed in the proximal colon, potentially through inflammatory or metabolic signaling pathways that alter mRNA synthesis or stability.

These findings have important therapeutic implications. In T1D, strategies aimed at restoring SNX17 expression or enhancing SNX17–NHE3 interactions could represent new avenues for preventing or mitigating fluid loss. In T2D, interventions that preserve NHE3 transcription, such as anti-inflammatory treatments or metabolic modulators, may be more effective. However, identifying effective targets will require further mechanistic studies to clarify the basis of NHE3 transcriptional dysregulation. Moreover, our findings that NHE3 is regulated by two endosomal sorting systems, SNX27 for trafficking and SNX17 for stability, broaden the landscape of potential molecular targets for intervention in intestinal diseases involving transporter dysfunction. In summary, this study not only elucidates a novel post-translational regulatory mechanism of NHE3 involving SNX17 but also highlights distinct and non-overlapping pathways by which T1D and T2D impair intestinal sodium absorption. These insights significantly advance our understanding of diabetic enteropathy and underscore the importance of tailoring therapeutic strategies to disease-specific molecular defects.

## Data available upon request from the senior author (VS)

### Grants

This study is partly supported by Cystic Fibrosis Foundation grant CF-SINGH20G0, National Institutes of Health grants R01-DK-116352, R24 DK099803, and P30-DK-89502 (the Hopkins Basic and Translational Research Digestive Diseases Research Core Center).

### Disclosures

The Authors declare no competing interests.

### Author Contributions

Conceived and Designed Research: VS; Performed Experiments: RL, LS, JY, RS; Analyzed Data: VS, RL, LS, JY; Interpreted Results: VS, MD; Prepared Figures: VS, RL, LS, JY, RS; Drafted Manuscript: VS; Edited and Revised Manuscript: VS, MD; Funding: VS, MD

Approved Final Manuscript: all authors

